# Xpert Ultra can unambiguously identify specific rifampicin resistance-conferring mutations

**DOI:** 10.1101/310094

**Authors:** Kamela C. S. Ng, Armand van Deun, Conor J. Meehan, Gabriela Torrea, Michèle Driesen, Siemon Gabriëls, Leen Rigouts, Emmanuel André, Bouke C. de Jong

## Introduction

The deluge of data produced by XpertMTB/RIF (Cepheid) can help improve global rifampicin-resistant tuberculosis (RR-TB) control strategies through molecular epidemiological surveillance (1, 2). Recently, a new version of the test – Xpert Ultra (hereinafter called Ultra) was released (3). Determining the relationship between RR-conferring *rpoB* mutations, Ultra probes, and melting temperature shifts (ΔTm) – the difference between mutant and wildtype melting temperatures – allows Ultra results to be utilized for rapid detection of RR-TB strains and related underlying *rpoB* mutations.

## Methods

To validate the usefulness of Ultra results for predicting specific mutations, we analyzed 10 RS-TB and 107 RR-TB strains from the Belgian Coordinated Collections of

## Microorganisms

in the Institute of Tropical Medicine, Antwerp, Belgium. These strains harbor 36 unique RR-conferring mutations determined by *rpoB* sequencing.

## Results

Overall, 31/32 (97%) mutations inside the Rifampicin Resistance Determining Region (RRDR) were correctly identified by Ultra. Of concern, mutation His445Arg gave a “RIF Resistance INDERTERMINATE” result among 3/4 strains tested while it was reported as RR in the initial validation study (3). The silent mutation Thr444Thr was not reported as RR (Figure 1). The RR-conferring mutations on codons 170, 250, 299, 482, and 491 situated outside the RRDR were not detected.

**Figure 1.**
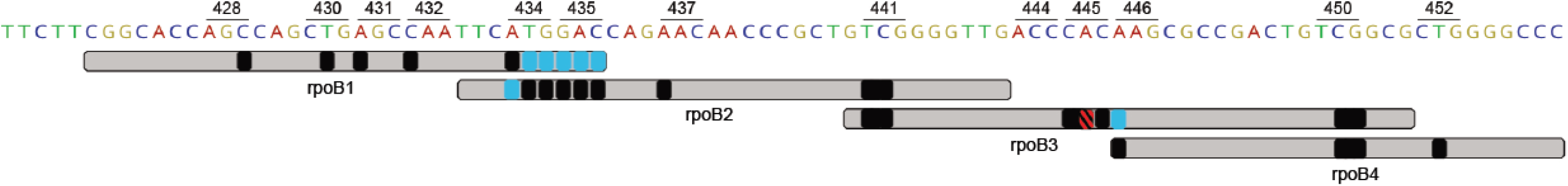
Overview of Xpert Ultra test results. The observed probe reactions for each RRDR mutation were overlaid on claimed probe coverage (light gray). Shown in black are probe reactions concordant with manufacturer claims; in blue are probe reactions missed by one probe but captured by another probe; and in red is a probe reaction representing “RIF Resistance INDETERMINATE” result on majority of strains tested. Results in striped pattern were superimposed for greater visibility.

The probe reactions observed were largely in agreement with previous results (3) albeit we noted that mutations Met434Val, Met434Thr and those in codon 435 were captured only by probe rpoB2; Ser450Leu and Ser450Trp were captured by both probes rpoB3 and rpoB4a, His445Arg was captured only by probe rpoB3; and Lys446Gln was captured only by probe rpoB4.

All mutations except those in codon 450 were associated with a negative ΔTm (Figure 2). The combination of ΔTm values with the capturing probes enabled to differentiate mutations in codons 428, 430, 431, 432, 434, 435, 441, 445, 446, and 452, including disputed mutations (4) (Table 1). Mutation Asp435Tyr was unambiguously distinguished from Asp435Val through probe rpoB2|ΔTm|, while mutations Ser441Gln and Ser441Leu were discriminated from the rest by|ΔTm| values of probes rpoB2 and rpoB3. Mutations His445Asp and His445Tyr were distinguished from disputed mutations His445Leu and His445Asn through probe rpoB3 |ΔTm|. Ser450Leu was distinguished from Ser450Trp by probe rpoB4A |ΔTm| except for one strain with an outlier rpoB4A Tm of 70.9°C in contrast to the other 13 strains with rpoB4A Tm of 73.3-73.8°C. The indeterminate result associated with His445Arg may be caused by its |ΔTm|=1.8°C compared with |ΔTm| typically exceeding 2°C for other mutations.

**Table 1.**
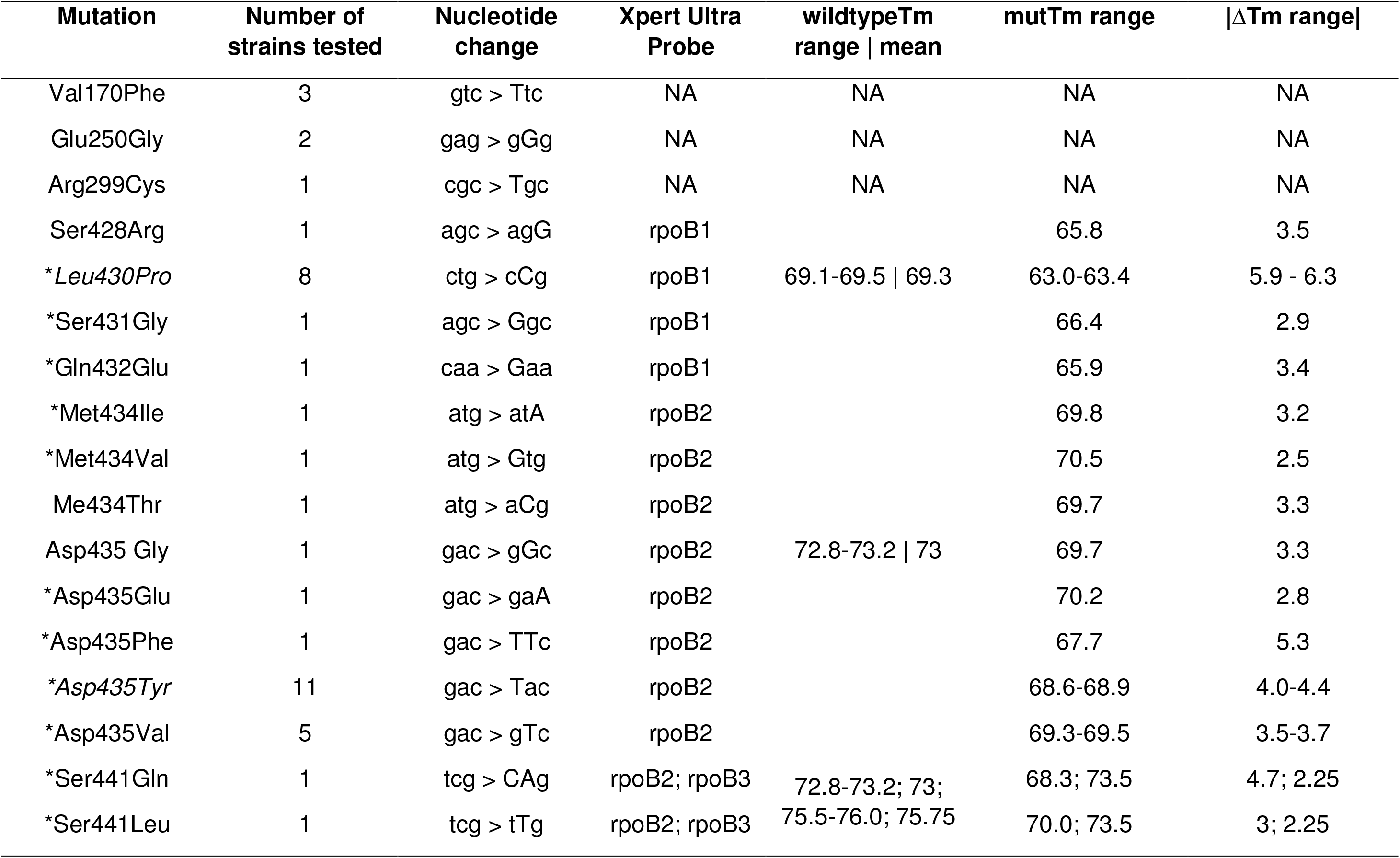
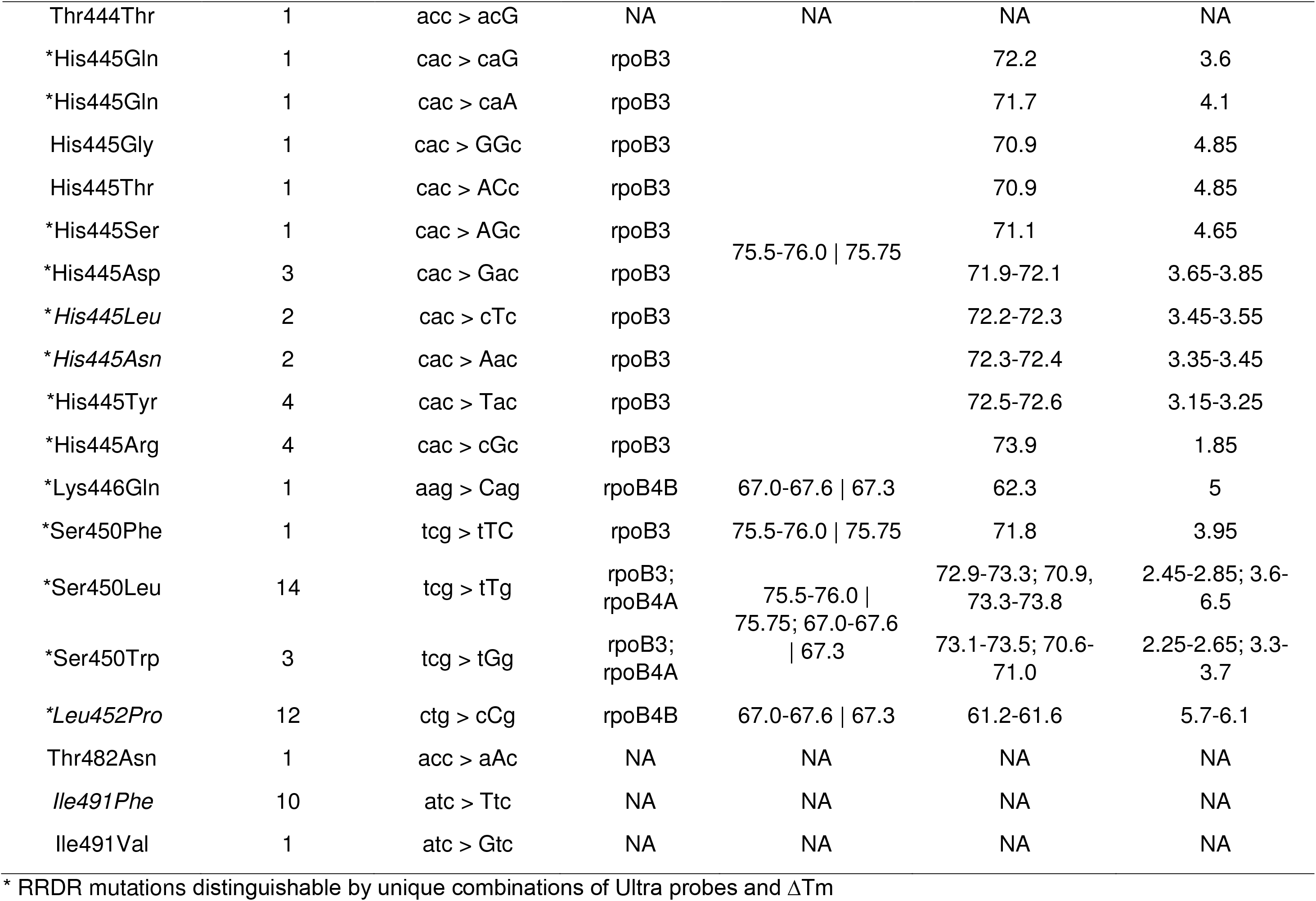
Xpert Ultra raw results – capturing probe, wildtype melt peak temperature (Tm) range and mean, mutant (mut) Tm range, and absolute value of melting temperature shift (ΔTm, range for multiple strains tested) – associated with rifampicin resistance-conferring mutations and corresponding nucleotide changes as determined by *rpoB* sequencing. Unique combinations of Ultra probe and ΔTm unambiguously differentiate most but not all mutations within the Rifampicin Resistance Determining Region (RRDR), including disputed ones in italics.

**Figure 2.**
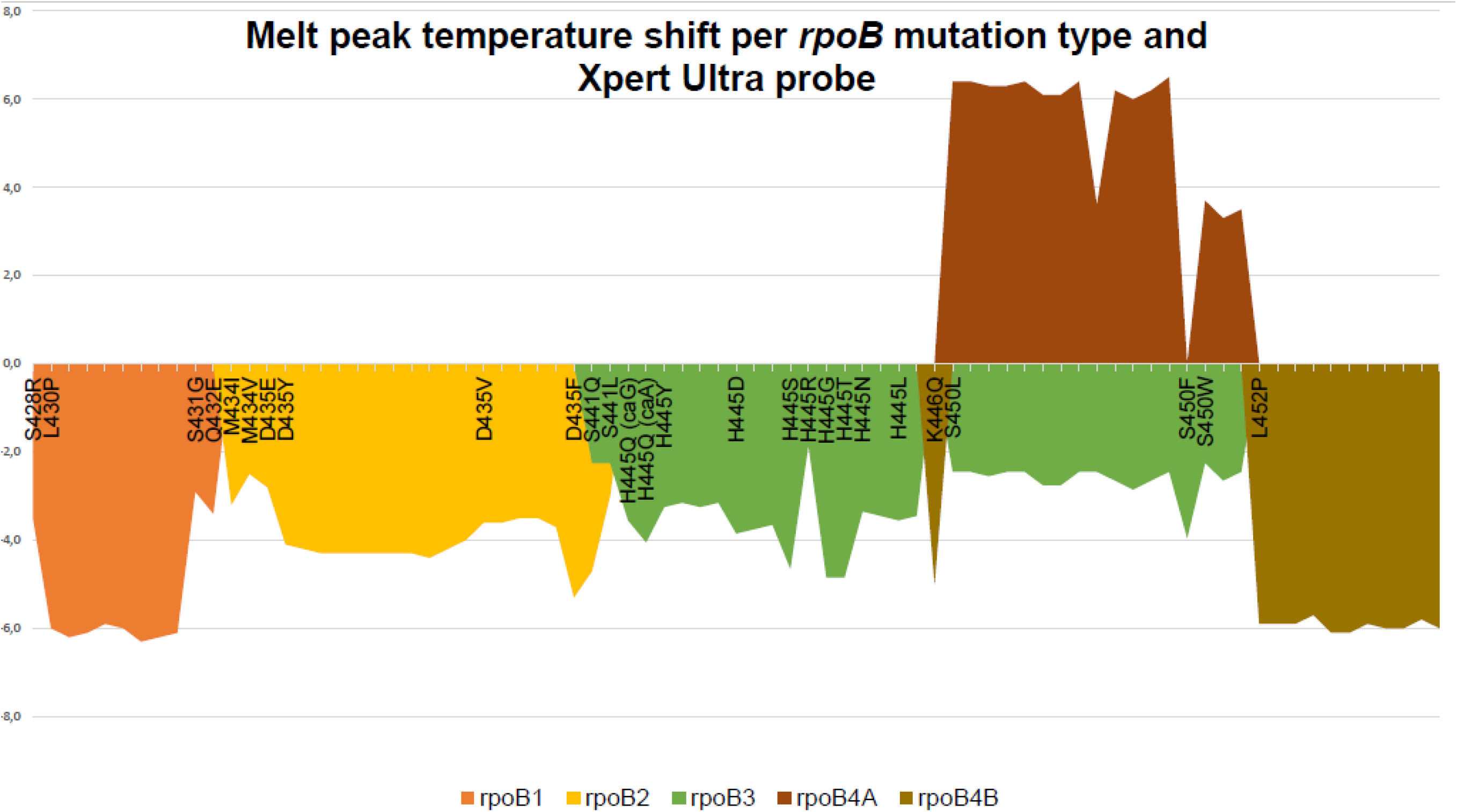
Melt peak temperature differences between Xpert Ultra probe-mutant and Xpert Ultra probe-wildtype amplicon hybrids per RRDR mutation type and probe. X-axis reflects the *rpoB* codon positions. The region representing isolates with the same mutation is marked by left-most mutation label.

## Conclusions

Our findings confirm the ability of Ultra to unambiguously identify a wide range of RRDR mutations. With the unprecedented roll-out of XpertMTB/RIF and associated connectivity solutions, such as DataToCare (Savics, Belgium) and GXAlert (SystemOne, USA) (2), Ultra results may be exploited to rule-out transmission between RR-TB patients in a specific setting, distinguish relapse from reinfection, and resolve discordance between an RR Ultra result and a low-level RS phenotypic result due to a disputed mutation. For such applications, it is key that ΔTm values are included in the exported results.

## Acknowledgement

This work was supported by Erasmus Mundus Joint Doctorate Fellowship grant 2016-1346 to KCSN.

